## Supplementary Figures for "Transcriptional Dynamics Uncover the Role of BNIP3 in Mitophagy during Muscle Remodeling in *Drosophila*"

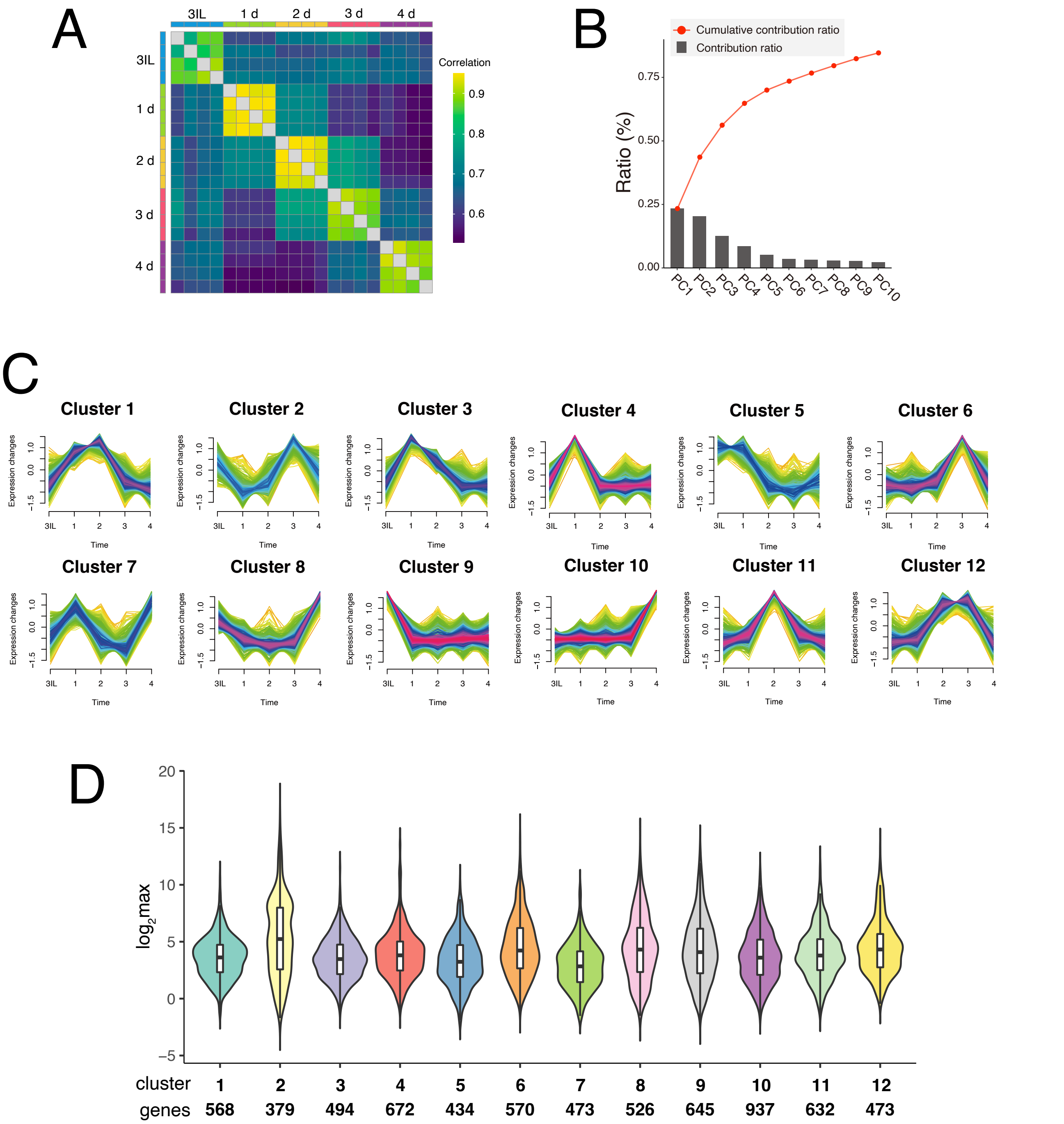


**Figure 1—figure supplement-1. RNA-seq of the DIOM remodeling**

(A) Heatmap of comparisons for each sample based on correlation coefficients. (B) The contribution ratios (bar) and cumulative contribution ratios (red-colored line) of PC1 to 10 are shown. (C) Fuzzy c-means clustering categorized expressed genes in DIOMs into twelve groups. (D) Violin plot showing the expression level of each cluster, with the number of genes in each cluster indicated at the bottom.

### Figure 1- fig supplement 1

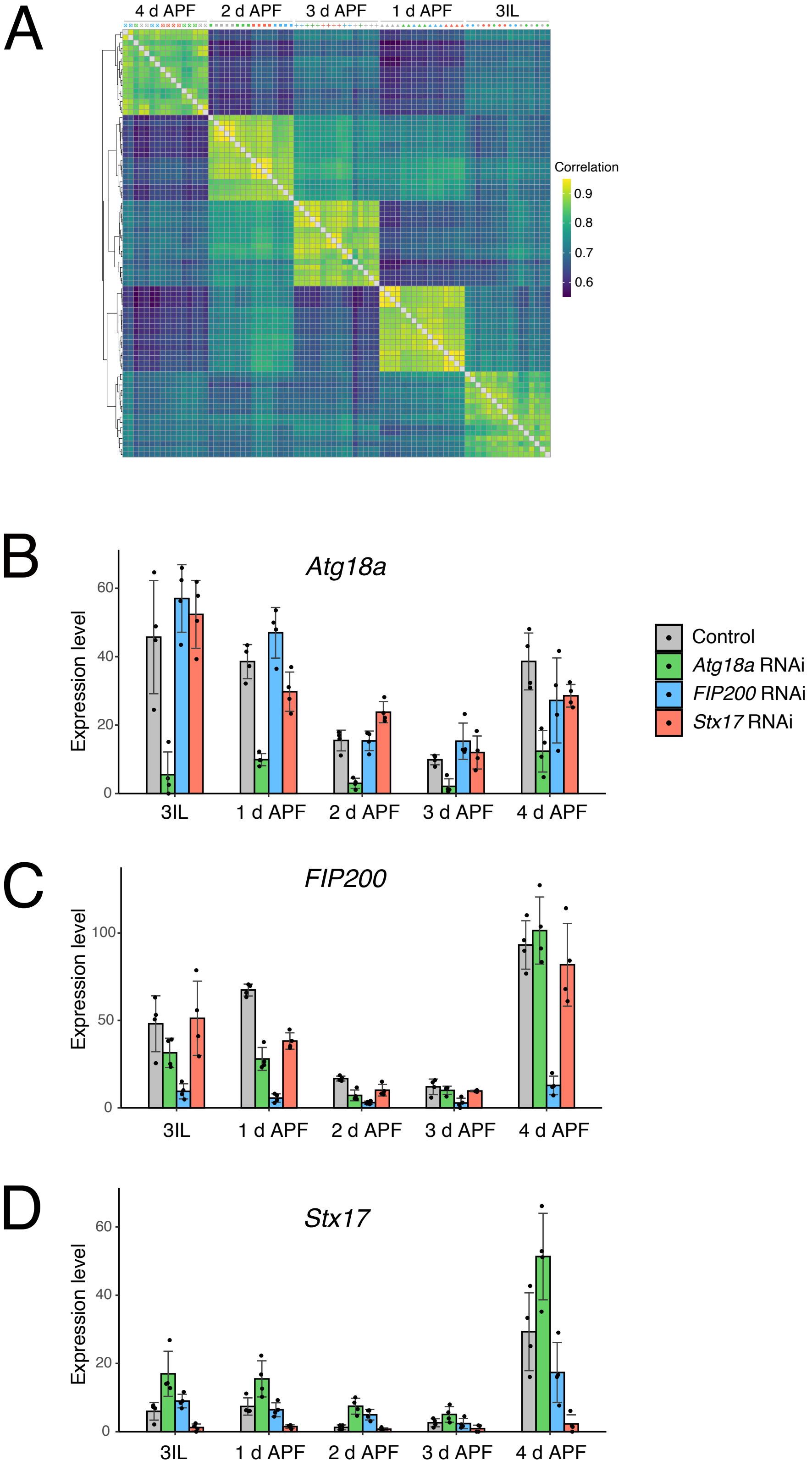


**Figure 2—figure supplement-1. Expression level of *Atg18a*, *FIP200*, and *Stx17* during DIOM remodeling** (A) Heatmap of comparisons for each sample based on correlation coefficients. (B-D) Expression levels of *Atg18a* (B), *FIP200* (C), and *Stx17* (D) in DIOMs under the indicated conditions. N=4.

**Figure 2- fig supplement 1**

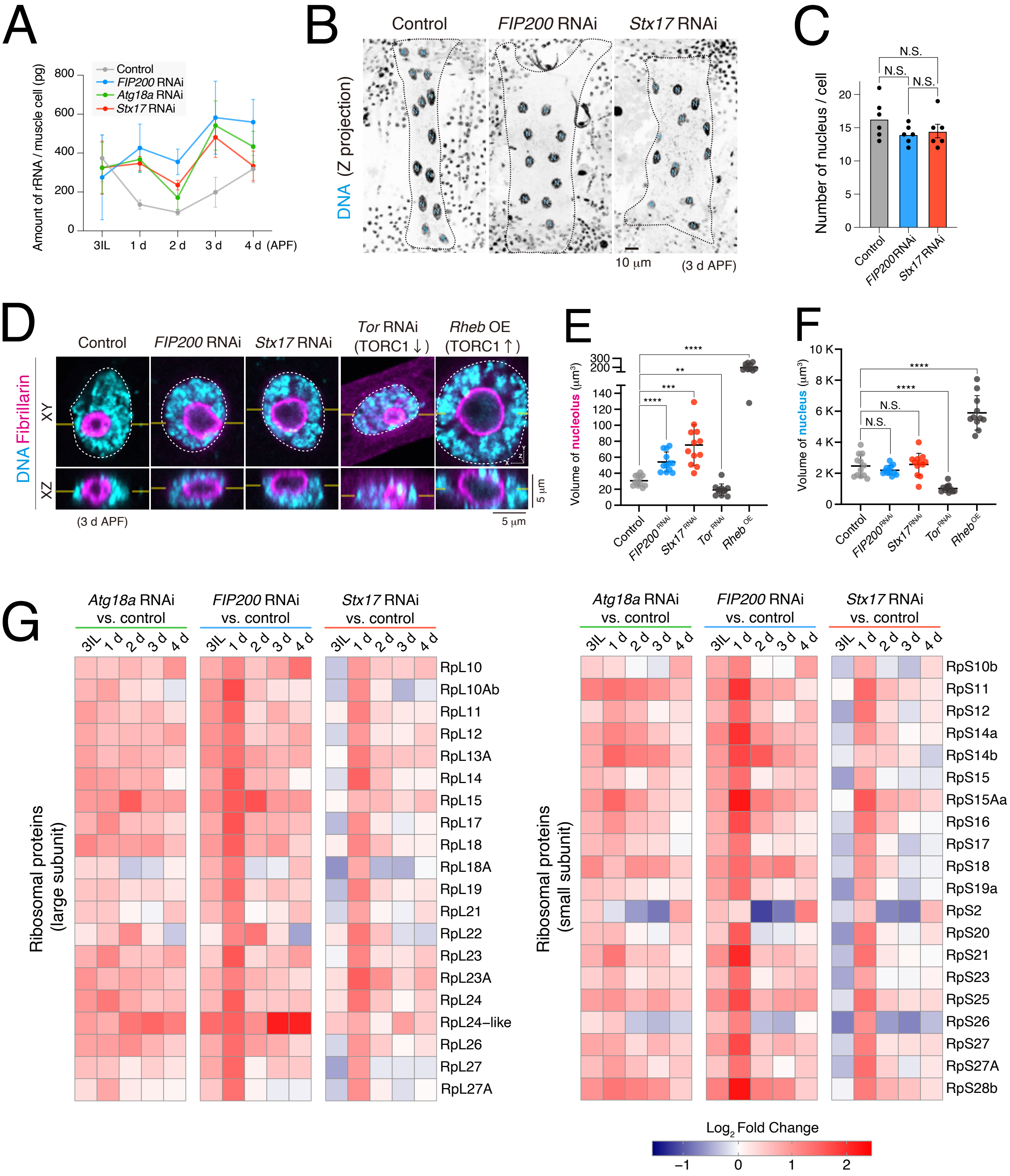


**Figure 2—figure supplement-2. Effect of autophagy deficiency on transcription of ribosomal RNA and protein**

(A) Time-course analysis of the amount of rRNA during DIOM remodeling in the indicated genotypes. (B and C) Loss of autophagy on the number of nuclei in DIOMs at 3 d APF. Nuclei in DIOMs were stained with Hoechst (B). N=6 (Sidak’ s test) (C). (D-F) Loss of autophagy on nucleolus volume in DIOMs at 3 d APF. DNA and fibrillarin were stained. XY and XZ planes are shown (D). The nucleolus (E) and nucleus (F) volume were calculated from Z-series images. N=12 (Dunnett’ s T3 multiple comparisons test). (G) A heatmap of the expression levels of RpLs (left) and RpSs (right) relative to control at each time point.

### Figure 2- fig supplement 2

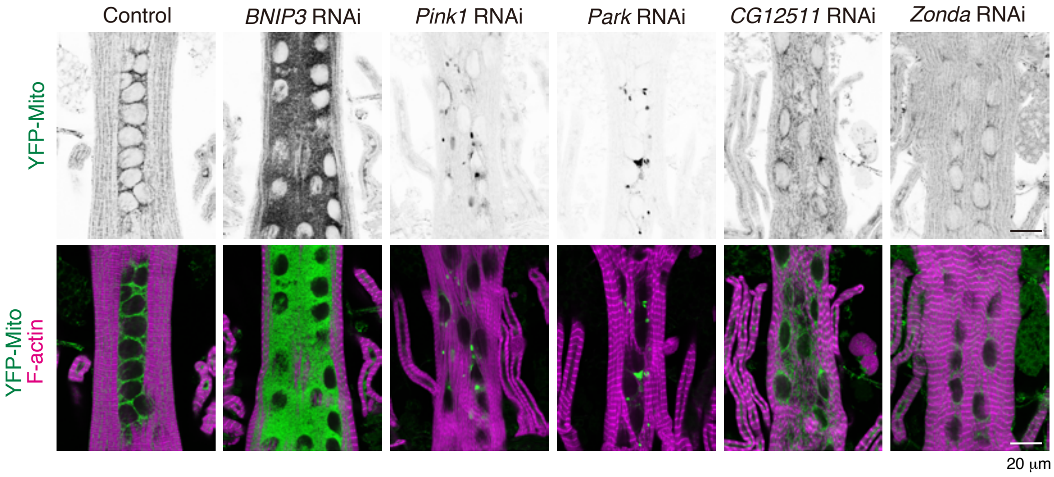


**Figure 3—figure supplement-1. Loss of BNIP3 on mitochondria and myofibrils in DIOMs**
YFP-Mito and Phalloidin-Alexa633 (F-actin) staining in DIOMs at 4 d APF in the indicated genotypes.

### Figure 3- fig supplement 1

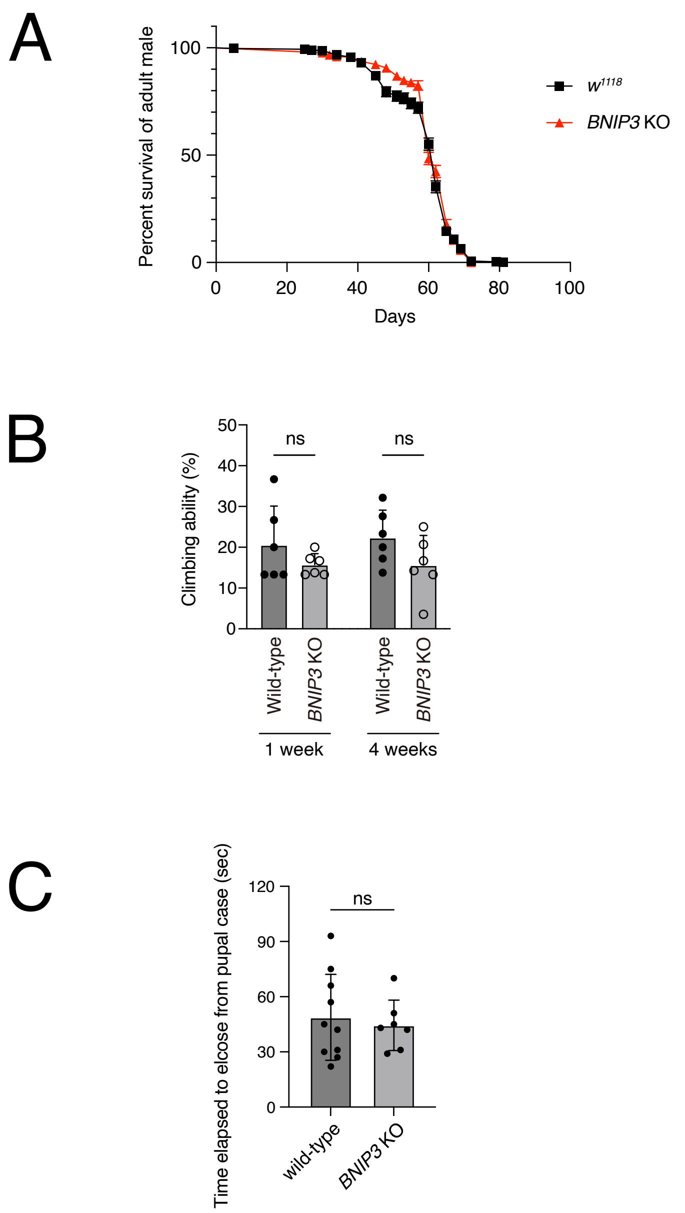


**Figure 3—figure supplement-2. Loss of BNIP3 on adult fly lifespan and mobility**
(A) Survival rates of wild-type and BNIP3 KO adult males. WT, N = 298; BNIP3 KO, N = 297. (Log-rank test). (B) Climbing ability of wild-type and BNIP3 KO adult males at 1 and 4 weeks old. N = 6 (Mann-Whitney test). (C) Elapsed time to eclosion in wild-type males and BNIP3 KO males. WT, N = 10; BNIP3 KO, N = 7 (Mann-Whitney test).

### Figure 3- fig supplement 2

A

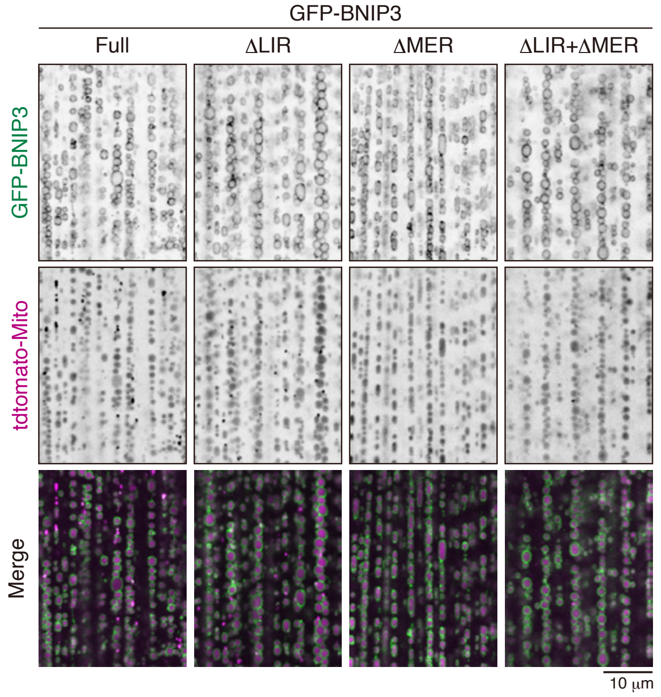


B

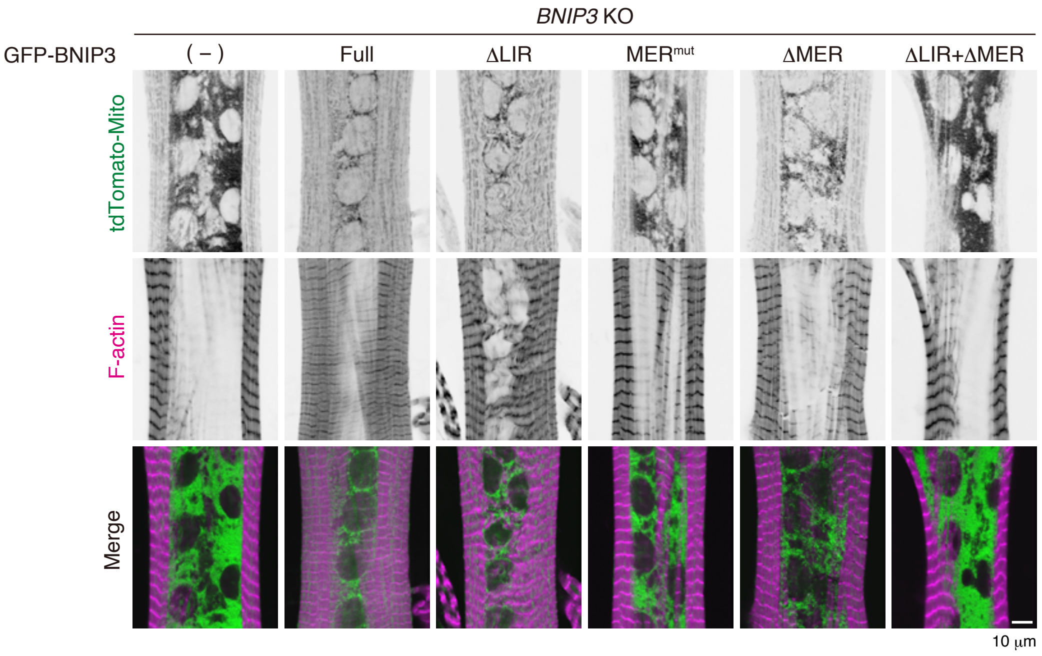


**Figure 5—figure supplement-1. GFP-BNIP3 constructs localize to the outer mitochondrial membrane** (A) Localization of GFP-BNIP3 constructs and mitochondria matrix-targeted tdTomato (tdTomato-Mito) in 3IL muscles. (B) BNIP3 rescue experiment in DIOMs at 4 d APF. The indicated GFP-tagged BNIP3 constructs and tdTomato-Mito were co-expressed in *BNIP3* KO flies using the GAL4/UAS system. Individual fluorescence channels are displayed separately.

### Figure 5- fig supplement 1

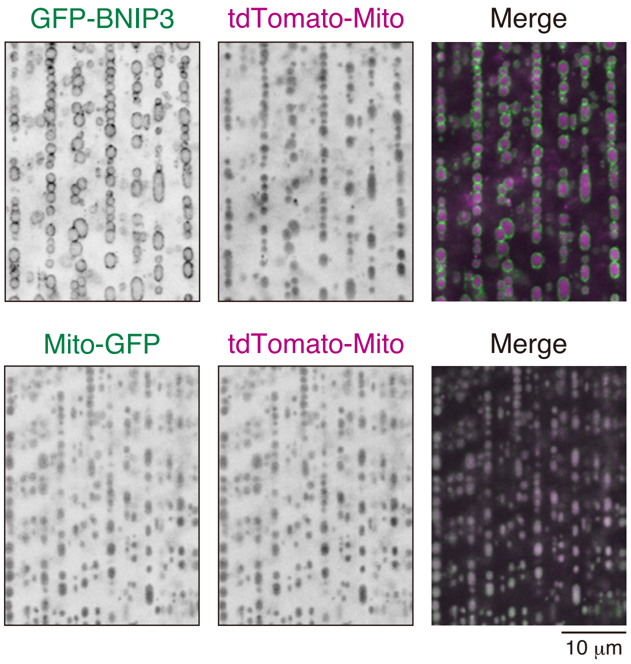


**Figure 5—figure supplement-2. Effect of GFP-BNIP3 overexpression on mitophagy flux in 3IL BWMs**
GFP-BNIP3 or mitochondria matrix-targeted GFP (Mito-GFP) were overexpressed together with tdTomato-Mito in 3IL BWMs. No clear difference in tdTomato-Mito signal was observed between samples with or without GFP-BNIP3 overexpression.

### Figure 5- fig supplement 2

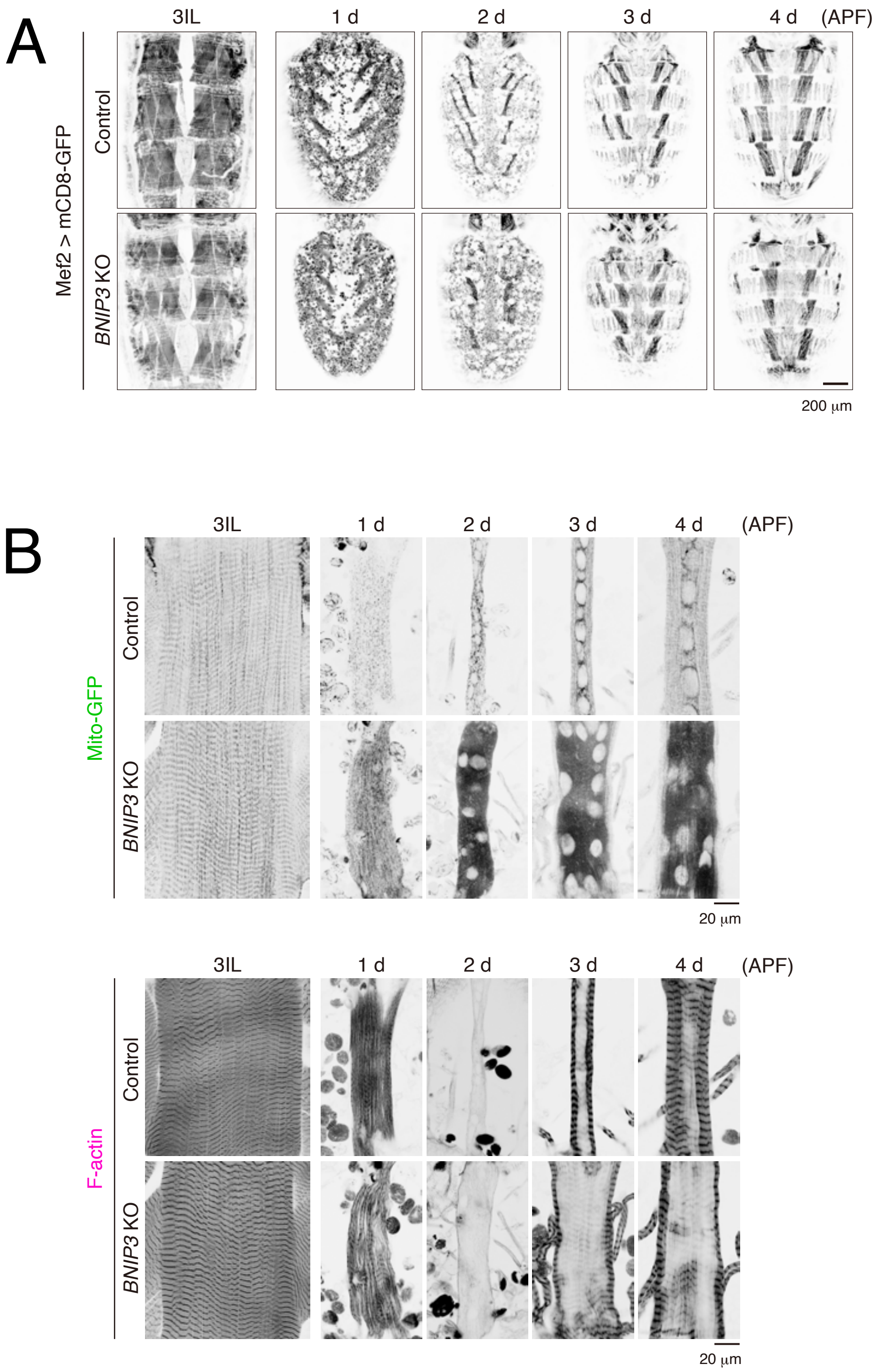


**Figure 6—figure supplement-1. The impact of *BNIP3* knockout on the shape of the DIOM during metamorphosis**

(A) Time-course microscopy of mCD8-GFP in dorsal muscles imaged through the cuticle in control and *BNIP3* KO animals from 3IL to 4 d APF. (B) Time course microscopy of Mito-GFP and F-actin in control or *BNIP3* KO during DIOM remodeling. Individual fluorescence channels are displayed separately.

### Figure 6- fig supplement 1
